# Improving rigor and reproducibility in western blot experiments with the blotRig analysis software

**DOI:** 10.1101/2023.08.02.551674

**Authors:** Cleopa Omondi, Austin Chou, Kenneth A. Fond, Kazuhito Morioka, Nadine R. Joseph, Jeffrey A. Sacramento, Emma Iorio, Abel Torres-Espin, Hannah L. Radabaugh, Jacob A. Davis, Jason H. Gumbel, J. Russell Huie, Adam R. Ferguson

## Abstract

Western blot is a popular biomolecular analysis method for measuring the relative quantities of independent proteins in complex biological samples. However, variability in quantitative western blot data analysis poses a challenge in designing reproducible experiments. The lack of rigorous quantitative approaches in current western blot statistical methodology may result in irreproducible inferences. Here we describe best practices for the design and analysis of western blot experiments, with examples and demonstrations of how different analytical approaches can lead to widely varying outcomes. To facilitate best practices, we have developed the blotRig tool for designing and analyzing western blot experiments to improve their rigor and reproducibility. The blotRig application includes functions for counterbalancing experimental design by lane position, batch management across gels, and analytics with covariates and random effects.

## Introduction

Proteomic technologies such as protein measurement with folin phenol reagent was introduced by Lowry et al in 1951(1). The resulting qualitative data are typically confirmed by a second, independent method such as western blot (WB)(2, 3). The WB method, first described by Towbin et al(4) and Burnette(5) in 1979 and 1981, respectively, uses specific antibody-antigen interactions to confirm the protein present in the sample mixture. Quantitative WB (qWB assay) is a technique to measure protein concentrations in biological samples with four main steps: (1) protein separation by size, (2) protein transfer to a solid support, (3) marking a target protein using proper primary and secondary antibodies for visualization, and (4) *semi*-quantitative analysis(6). Importantly, qWB data is considered *semi*-quantitative because methods to control for experimental variability ultimately yield relative comparisons of protein levels rather than absolute protein concentrations(2, 3, 7, 8). Current methodologies do not sufficiently account for the diverse sources of variability, producing highly variable results between different laboratories and even within the same lab(9–11) . Indeed, qWB data exhibits more variability compared to other experimental techniques such as enzyme linked immunosorbent assay (ELISA)(12). For example, results have shown that qWB can produce significant variability in detecting host cell proteins and lead to researchers missing or overestimating true biological effects(13). This in turn results in publication of irreproducible qWB interpretations, which leads to loss of its credibility(11). In the serious cases, qWB results may even provide clinical misdiagnosis(14) that could impact on a larger public health concern due to the prevalence of WB in biomedical research, such as diagnosis of SARS-CoV2 infection(15).

The process of recognizing and accounting for variability in WB analyses will ultimately improve reproducibility between experiments. A growing body of studies has shown that this requires a fundamental shift in the experimental methodology across data acquisition, analysis, and interpretation to achieve precise and accurate results(2, 9–11). Here we highlight experimental design and practices that enable a statistics-driven approach to improve the reproducibility of qWBs. Specifically, we discuss major sources of variability in qWB including the non-linearity in antibody signal(2); imbalanced experimental design(11); lack of standardization in the treatment of technical replicates(2, 16); and variability between protein loading, lanes, and blots(3, 8, 17). To address these issues, we provide new comprehensive suggestions for quantitative evaluation of protein expression by combining linear range characterization for antibodies, appropriate counterbalancing during gel loading, running technical replicates across multiple gels, and by taking careful consideration on the analysis method. By applying these experimental practices, we can then account for more sources of variability by running analysis of covariance (ANCOVA) or generalized linear mixed models (LMM). Such approaches have been shown to successfully improve reproducibility compared to other methods(11).

To help improve WB rigor we developed the blotRig protocol and application harnessing a database of 6000+ western blots from N = 281 subjects (rats and mice) collected by multiple UCSF labs on core equipment. To demonstrate blotRig best practices in a real-world experiment, we carried out prospective multiplexed WB analysis of protein lysate from lumbar cord in rodent models of spinal cord injury (SCI) (N=29 rats) in 2 groups (experimental group & control group). In order to show that these experimental suggestions could improve qWB reproducibility, we compared different statistical approaches to handling loading controls and technical replicates. Specifically, we applied two strategies to integrate loading controls: (i) normalizing the target protein levels by dividing by the loading control or (ii) treating the loading control as a covariate in a LMM. Additionally, we analyzed technical replicates in four ways: 1) assume each sample was only run once without replication, 2) treat each technical replicate as an independent sample, 3) use the mean of the three technical replicate values, and 4) treat the replicate as a random effect in a LMM. Altogether, we found that the statistical power of the experiment was significantly increased when we used loading control as a covariate with technical replicates as a random effect during analysis. In addition, the effect size was increased, and the p-value of our analysis decreased when using this LMM, suggesting the potential for greater sensitivity in our WB experiment when using this approach(18). Through rigorous experimental design and statistical analysis we show that we can account for greater variability in the data and more clearly identify underlying biological effects.

## Materials and Methods

### Animals

All experiments protocol were approved by the University Laboratory Animal Care Committee at University of California, San Francisco (UCSF, CA, USA) and followed the animal guidelines of the National Institutes of Health Guide for the Care and Use of Laboratory animals(19). Male Simonsen Long Evans rats (188-385 g; Gilroy (Santa Clara, CA, USA), (N=29) aged 3 weeks were housed under standard conditions with a 12-h light–dark cycle (6:30 am to 6:30 pm) and were given food and water ad libitum. The animals were housed mostly in pairs in 30 × 30 × 19-cm isolator cages with solid floors covered with a 3 cm layer of wood chip bedding. The experimenters were blind to the identity of treatments and experimental conditions, and all experiments were designed to minimize suffering and limit the number of animals required.

### Experimental methodology

In accordance with established quality standards for preclinical neurological research(20), experimenters were kept blind to experimental group conditions throughout the entire study. Western blot loading order was determined a *priori* by a third-party coder, who ensured that a representative sample from each condition was included on each gel in a randomized block design. The number of subjects per condition was kept consistent across groups for each experiment to ensure that proper counterbalancing could be achieved across independent western runs. All representative western images presented in the figures represent lanes from the same gel. Sometimes, the analytical comparisons of interest were not available on adjacent lanes even though they come from the same gel because of our randomized counterbalancing procedure.

### Western blot

The example western blot data used in this paper are taken from a model of spared nerve injury in animals with spinal cord injury. The nerve injury model used is based on models from pain literature(21), where two of the three branches of the sciatic nerve are transected, sparing the sural nerve (SNI)(22). Two surgeons perform the procedure simultaneously, with injuries occurring 5 minutes apart. The spinal cord of animals was obtained based on fluid expulsion model(23) and a 1 cm section of the lumbar region was excised at the lumbar enlargement section. The tissue was then preserved in a -80 degree freezer until it was needed for an experiment, at which point it was thawed and used to run a Western blot. We conducted a Western blot analysis on 29 samples from animals using standard biochemical methods. We measured the protein levels of the AMPA receptor subunit GluA2 and used beta-actin as a loading control. The data from these experiments was then aggregated and used for statistical analysis. Our WB protocol:

#### 1. Protein Assay

We assayed sample protein concentration using a bicinchoninic acid (BCA assay (Pierce) for reliable quantification of total protein using a plate reader (Tecan; GeNios) with triplicate samples (technical replicates) detected against a Bradford Assay (BSA) standard curve. Technical replicates are multiple measurements that are performed under the same conditions in order to quantify and correct for technical variability and improve the accuracy and precision of the results (48). We run the same WB loading scheme three times (technical replicates of the entire gel) and measured the protein levels of AMPA receptors.

#### 2. Polyacrylamide Gel Electrophoresis and Multiplexed Near-Infrared Immunoblotting

The approach involved performing serial 1:2 dilutions with cold Laemmli sample buffer in room temperature; 15 μg of total protein per sample was loaded into separate lanes on a precast 10–20% electrophoresis gel (Tris-HCl polyacrylamide, BioRad). The blotRig software helps counterbalance sample positions across the gel by treatment condition., (**Fig 2**). A kaleidoscope ladder was loaded on the first lane of each gel to confirm molecular weight (**Fig 2**). The gel was electrophoresed for 30 minutes at 200 V in SDS buffer (25 mm Tris, 192 mm glycine, 0.1% SDS, pH 8.3; BioRad). Protein was transferred to a nitrocellulose membrane in cold transfer buffer (25 mm Tris, 192 mm glycine, 20% ethanol, pH 8.3). Membrane transfer was confirmed using Ponseau stain followed by a quick rinse and blocking in Odyssey blocking buffer (Li-Cor) containing Tween-20.

The membrane was blocked for 1 h in Odyssey Blocking Buffer (Li-Cor) containing 0.1% Tween-20, followed by an overnight incubation in primary antibody solution at 4°C. Membrane incubation was done in a primary antibody solution containing Odyssey blocking buffer, Tween-20, appropriate primary antibody receptor targeting1:2,000 mouse PSD-95 (cat # MA1-046,Thermofisher), 1:200 rabbit GluA1 (cat # AB1504, Millipore), 1:200 rabbit GluA2 (cat # AB1766, Millipore), 1:200 rabbit pS831(cat # 04-823, Millipore), 1:200 p880 (cat#07-294, Millipore) or 1:1,500 mouse actin loading control (cat # 612857, BD Transduction)]. Following incubation, the membrane was washed 4 × 5 min with Tris-buffered saline containing 0.1% Tween 20 (TTBS) and incubated in fluorescent-labeled secondary antibody (1:30K LiCor IRdye appropriate goat anti-rabbit in Odyssey blocking buffer plus 0.2% Tween 20) for 1 h in the dark. Subsequent to 4 × 5 min washes in TTBS, followed by a 5 min wash in TBS.

Membrane incubation was used to detect the presence of a specific protein or antigen on a membrane. In this case, the membrane was incubated with a fluorescently labeled secondary antibody solution that was specifically tuned to the emission spectra of the laser lines used by the Li-Cor Odyssey quantitative near-infrared molecular imaging system instrument. This allows for specific detection of the protein of interest on the membrane. The sample is then imaged using an infrared imaging system that is optimized for detecting the specific wavelengths of light emitted by the fluorescent label. Additional rounds of incubation and imaging are performed to detect additional proteins using the multiplexing functionality of the Li-Cor instrument, with each round adding new bands at different molecular weight ranges. This allows for the detection of multiple proteins in the same sample, maximizing the proteomic detection.

#### 3. Quantitative Near-IR Densitometric Analysis

Using techniques optimized in the Ferguson lab(24, 25), we established near-infrared labeling and detection techniques (Odyssey Infrared Imaging System, Li-Cor) to quantify linear intensity detection of fluorescently labeled protein bands. The biochemistry is performed in a blinded, counterbalanced fashion, and three independent replications of the assay are run on different days(26). Fluorescent Western blotting utilizes fluorescent-labeled secondary antibodies to detect the target protein, which allows for more sensitive and specific detection compared to chemiluminescence(9, 27, 28). Additionally, fluorescence imaging allows multiple detection of a target protein and internal loading control in the same blot, which enables more accurate correction of sample-to-sample and lane-to-lane variation(9, 29, 30). This provides a more accurate and reliable quantification of the target protein, making it a popular choice for quantitative analysis of WB data.

#### 4. Blinding

It is good practice for the pipetting experimenter to remain blind to experimental conditions during gel loading, transfer, and densitometric quantification. We achieved this using de-identified tube codes and *a priori* gel loading sequences that were developed by an outside experimenter using the method implemented in the blotRig software.

### Statistical analyses

Statistical analyses were performed using the R statistical software (58,59). Our WB data was analyzed using parametric statistics. The WB was run using three independent replications and covariance corrected by beta-actin loading control, with replication statistically controlled as a random factor. Significance was assessed at p < 0.05(24, 25, 31, 32). We report estimated statistical power and standardized regression coefficient effect sizes in the results section.

All ANOVAs were run using the *stats* R package; standardized effect size was calculated using the *parameters* R package (58). Linear mixed models were run using the *lme4* R package. Observed power was calculated by Monte Carlo simulation (1000x) run on the fitted model (either ANOVA or LMM) using the *simR* package (59). For the development of the blotRig interface, the R packages used included: *shiny, tidyverse, DT, shinythemes, shinyjs,* and *sortable* (58–64). You can access the blotRig analysis software, which includes code for inputting experimental parameters for all Western blot analysis, through the following link: https://github.com/ucsf-ferguson-lab/blotRig/

## Results

### DESIGNING REPRODUCIBLE WESTERN BLOT EXPERIMENTS

#### 1. Determining linear range for each primary antibody

Most WB analyses assume *semi*-quantitatively that the relationship between qWB assay optical density data (i.e. western band signal) and protein abundance is linear(2, 3, 9, 16). Accordingly, most qWB analyses use statistical tests that assume a linear effect. However, recent studies have shown that the relationship can potentially be highly non-linear(17) As **Figure 1** illustrates, the WB band signal can become non-linearly correlated with protein concentrations at low and high values. This may result in inaccurate quantification of relative target protein amount in the experiment and violates the assumptions for linear model which can lead to false inferences. To address the assumption of linearity, it is important to first determine the optimal linear range for each protein of interest so that one can be confident that a unit change in band density reflects a linear change in protein concentration. This enables an experimenter to accurately quantify the protein of interest and apply linear statistical methods appropriately for hypothesis testing.

**Figure 1.**
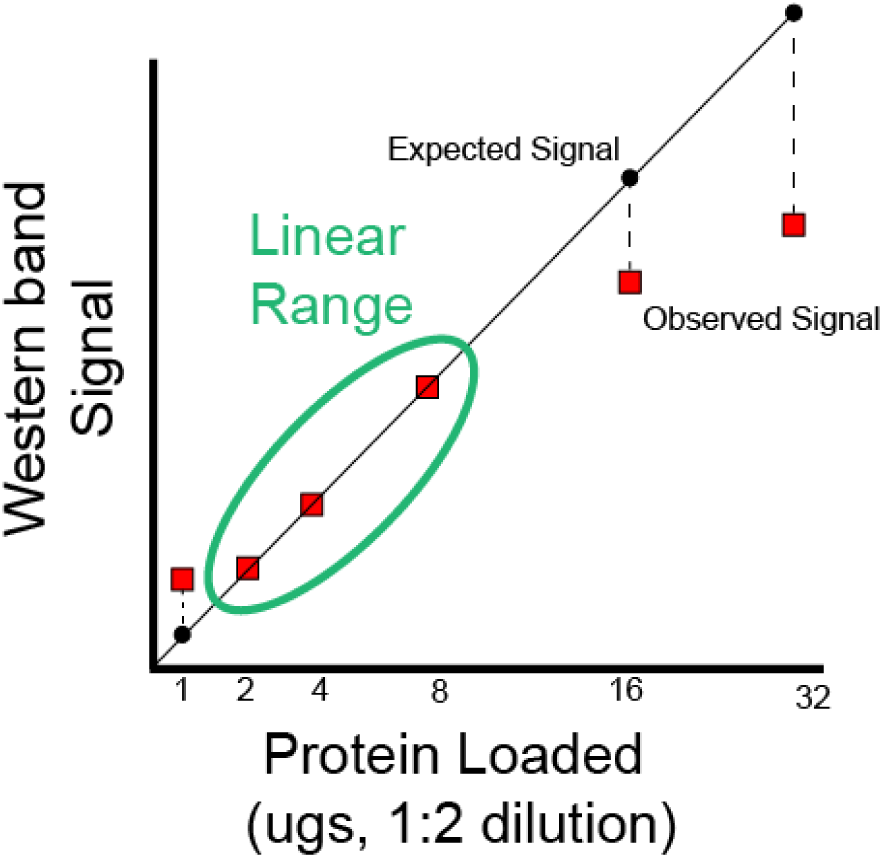
Determining linear range of antibodies to optimize parametric analysis of Western blot data. When small or large protein concentrations are loaded, there is often a possibility that their representation on western blot band density may become non-linear. If there is a disconnect between the observed and expected protein concentrations, results may be inaccurate. Thus determining the linear range wherein, a one-unit increase in protein is reflected in a linear increase in band density for each western blot antibody is a crucial initial step to ensure confidence in reproducibility of the linear models commonly applied to western blot data analysis.

**Figure 2.**
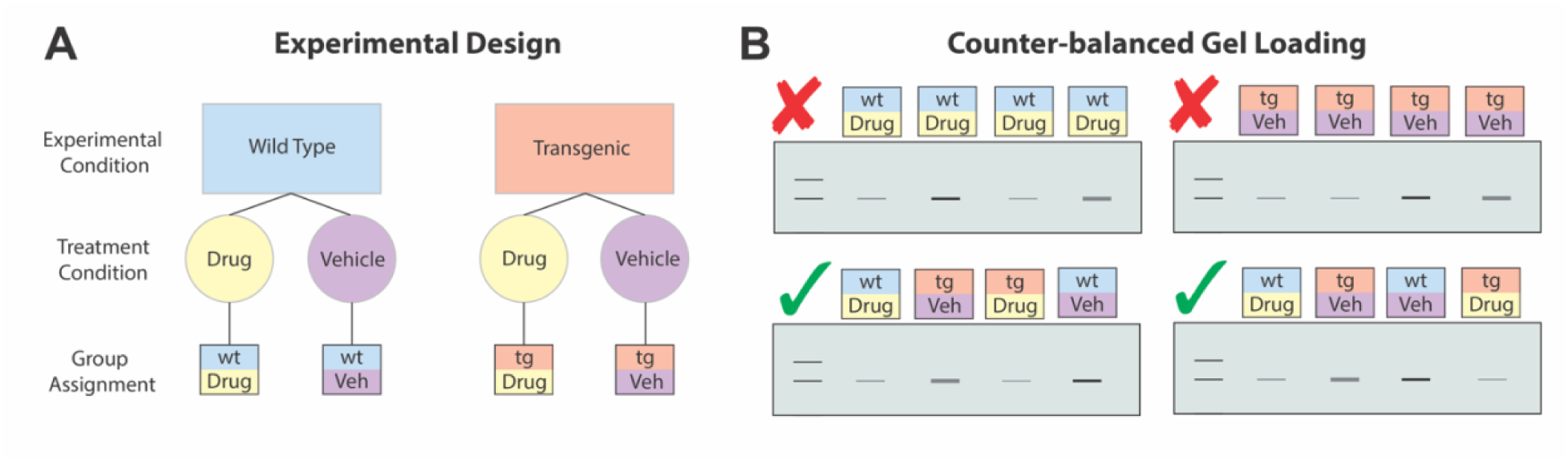
Counterbalancing to reduce bias. **A, Experimental design**. A simple hypothetical experimental design for illustrating counterbalancing. Two experimental groups (Wild Type vs Transgenic), with two treatments (Drug vs Vehicle) analyzed within each individual. This 2 (Experimental Condition) by 2 (Tissue Area) design yields four groups. **B, Counter-balanced Gel Loading**. The goal of appropriate counterbalancing is to optimize the sequence in which samples are loaded such that groups are represented equally across the gel. Those with red X have with the experimental groups and treatment condition grouped in the same area of the gel, and thus variability across the gel may be conflated with group differences. In contrast, those with the green check are organized so that experimental condition and treatment condition are better placed to reduce the possibility of any single group being over-represented in a particular area of the gel.

#### 2. Counterbalancing during experimental design

Counterbalancing is the practice of having each experimental condition represented on each gel and evenly distributing them to prevent overrepresentation of the same experimental groups in consecutive lanes. For example, imagine an experimental design in which we are studying two experimental groups (wild type and transgenic animals) and are also looking at two treatment conditions (Drug and Vehicle). This design gives us four groups total (Drug-treated Wild Type, Vehicle-treated Wild Type, Drug-Treated Transgenic and Vehicle-treated Transgenic) (**Figure 2A**). During WB gel loading, experimenters often distribute their samples unevenly such that certain experimental conditions may be missing on some gels or samples from the same experimental condition are loaded adjacently on a gel. This is problematic because we know that polyacrylamide gel electrophoresis (PAGE) gels are not perfectly uniform, reflecting a source of technical variability(34); in the worst case, if we have only loaded a single experimental group on a gel and found a significant effect of the group, we cannot conclude if the effect is due to the experimental condition or a technical problem of the gel. At minimum, experimenters should ensure that every group is represented on each gel to avoid confounding technical gel effects with experimental differences.

In addition, experimenters can further counter technical variability by arranging experimental groups on each gel to ensure adequately counterbalanced design assuming the uniformed protein concentration and fluid volume of all samples. This importantly addresses the variability due to physical effects within an individual gel. In our example, this means alternating the tissue areas and experimental conditions as much as possible to minimize similar samples from being loaded next to one another (**Figure 2B**). By spreading the possibility of technical variability across all samples by counterbalancing across and within gels, we can mitigate potential technical effects that can bias our results. Proper counterbalancing also enables us to implement more rigorous statistical analysis to account for and remove more technical variability(24, 25, 31, 32). Overall, this will help to ensure that experimenters can find the same result in the future and improve reproducibility.

#### 3. Technical Replication

Technical replicates are used to measure the precision of an assay or method by repeating the measurement of the same sample multiple times (11). The results of these replicates can then be used to calculate the variability and error of the assay or method(11). This is important to establish the reliability and accuracy of the results. Most experimenters acknowledge the importance of running technical replicates to avoid false positives and negatives due to technical error(11). Even beyond extreme results, technical replicates can account for the differences in gel makeup, human variability in gel loading, and potential procedural discrepancies. In fact, most studies run at least duplicates; however, the experimental implementation of replicates (e.g., running replicates on the same gel or separate gels) as well as the statistical analysis of replicates (e.g., dropping “odd-man-out” or taking the mean or standard deviation) can differ greatly(35, 36). This experimental variability ultimately impedes our ability to meaningfully compare results. For experimenters to establish accuracy and advance reproducibility in WB experiments, it is important to implement standardized and rigorous protocols to handle technical replicates(9, 11). In doing so, we can further reduce the technical variability with statistical methods during analysis.

As underscored previously, we recommend that technical replicates are counterbalanced on separate gels to mitigate any possible gel effect. Additionally, by running triplicates, we can treat replicates as a random effect in a LMM during statistical analysis. Importantly, triplicates provide more values to measure the distribution of technical variance to ensure the robustness of the LMM than only running duplicates. This approach isolates and removes technical variance from biological variation which ultimately improves our sensitivity for true experimental effects(37).

In the following demonstration of statistical methods, we replicated all WB analyses in triplicate with a randomized counterbalanced design. We then explore how the way in which technical replicates and loading controls are incorporated into analysis can have a significant impact on both the sensitivity of our results and the interpretation of the findings.

### STATISTICAL METHODOLOGY TO IMPROVE WESTERN BLOT ANALYSIS

#### 1. Loading control as a covariate

Most qWB assay studies use loading controls to ensure that there are no biases in total protein loaded in a particular lane(3, 9, 26). The most common way that loading controls are used to account for variability between lanes is by normalizing the target protein expression values by dividing it by the loading control values, resulting in a ratio between target protein to loading control(3, 38, 39). However, ratios may violate assumptions of common statistical test used to analyze qWB (e.g., t-test, ANOVA, etc.)(40) This ultimately hinders the ability to statistically account for the variance in qWB outcomes and have a reliable estimate of the statistics. We instead propose to include loading control values as a covariate – a variable that is not our experimental factors that may affect the outcome of interest and presents a source of variance that we wish to account for(41). For instance, we know the amount of protein loaded is a source of variability in WB quantification, so we can use the loading control as a covariate to adjust for that variance. In doing so, we extend the method of ANOVA into that of ANCOVA(42). This approach accounts for the technical variability present between lanes while meeting the necessary assumptions for parametric statistics which helps curb bias and averts false discoveries.

#### 2. Replication and Subject as a random effect

Most WB studies use ANOVA, a test that allows comparison of the means of three or more independent samples, for quantitative analysis of WB data(40). One of the assumptions in ANOVA is the independence of observations(40). This is problematic because we often collect multiple observations from the same analytical unit, for example different tissue samples from a single subject, or technical replicates. As a result, those observations don’t qualify as independent and would rather be analyzed using models controlling for variability within units of observations (e.g., the animal) to mitigate inferential errors (false positives and negatives)(43) caused by what is known as pseudoreplication. This arises when the quantity of measured values or data points surpasses the number of actual replicates, and the statistical analysis treats all data points as independent, resulting in their full contribution to the final result (65)

In addition, when conducting experiments, it is important to consider the randomness of the conditions being observed. Treating both subjects and conditions as fixed effects can lead to inaccurate p-values. Instead, subjects/ animals should be treated as random effects and the conditions should be considered as a sample from a larger population(44). This is especially important when collecting data from different replicates or gels, as the separate technical replicate runs should be considered as random.

In Figure 3 we used a simple experimental design comparing the difference in a target protein between two experimental groups to demonstrate four of the most common ways researchers tend to analyze western blot data: 1) running each sample once without replication, 2) treating each technical replicate as an independent sample, 3) taking the mean of technical replicate values, and 4) treating subject and replication as a random effect (**Figure 3**). We then tested how effect size, power, and p value are affected by each of these strategies to get a sense of how much these estimates vary between analyses. For each of these strategies, we also tested the difference between using the ratio of target protein to loading controls versus using loading control as a statistical covariate.

**Figure 3.**
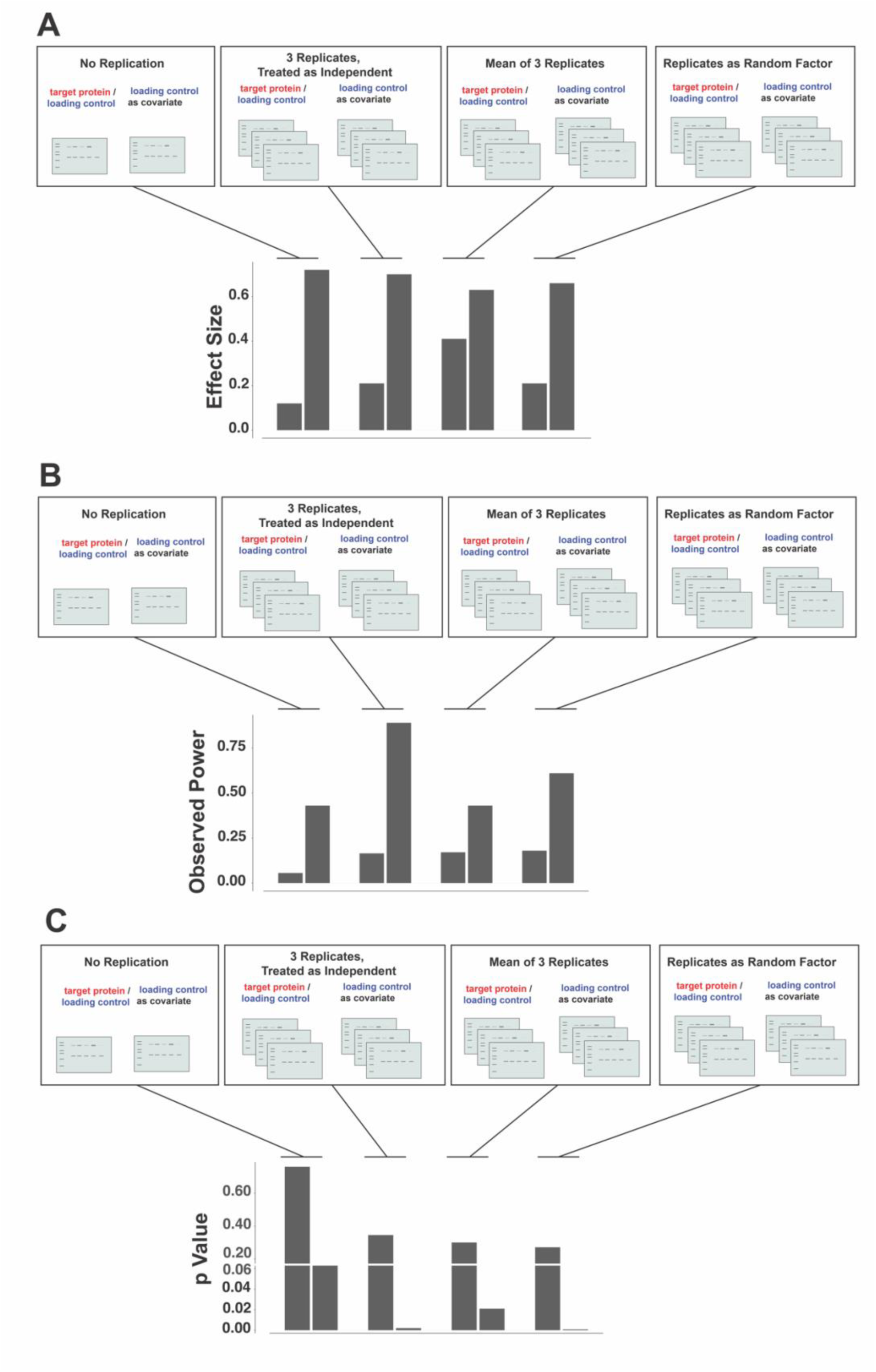
Effect of different replication and loading control strategies on statistical outcomes. Eight possible strategies are shown, representing the most common ways in which replication and loading controls are treated in a typical Western blot analysis. Four replication strategies: either no replication at all, 3 technical replicates treated as independent, mean of three replicates, or replicate treated as a random effect in a linear mixed model. These are crossed with two loading control strategies: either target protein is divided by loading control, or loading control is treated as a covariate in a linear mixed model. **A**, **Effect Size:** Standardized effect size is generally improved when loading control is treated as a covariate, compared to using a ratio of the target protein and loading control values. **B, Power:** By treating each replication as independent the power is increased (due to the i*naccurate* assumption that technical replicates are not related, thus artificially tripling the n). Conversely, including the variability inherent in technical replicates as a part of the statistical model, we work to identify and account for a major source of variability, thus improving power in a more appropriate way. **C, P value:** As expected, when each replication is inaccurately treated as independent the p value is low (due to artificially inflated n). We found that using the mean of replications and loading controls as covariates also resulted in a p value below 0.05. The smallest p value was found when including replication as a random factor. Across each of these statistical measures, only when replication is included as a random factor and loading control as a covariate do we see a strong effect size, high power, and low p value.

In the first scenario, we imagined that no technical replication was run at all (by using only the first replication). With this strategy, we found that standardized effect size is weak, power is low, and the p value was high (**Figure 3**). Second, we demonstrate how analytical output would be different if we did run three technical replicates, but treated each as independent. As discussed above, this strategy does not take into account the fact that each sample is being run three times, and consequently the overall n of your experiment is artificially tripled! As one might expect, observed power is quite high, and our p value is low (< 0.05). Power is increased by an increase in sample size, so it is not surprising that the power is much higher if we erroneously report that we have a 3X larger sample size (i.e., pseudoreplication) (65). In this case, the observed power is inflated and an artifact of inappropriate statistics, and the probability of a false positive is considerably increased with respect to the expected 5%.

So, what would be a more appropriate way to handle technical replicates? One method that researchers often use is to take the mean of their technical replicates. This does ensure that we are not artificially inflating our sample size, which is certainly an improvement over the previous strategy. With this strategy, we do find that our p value is less than 0.05 (when loading control is treated as a covariate). But we also see that our power is still low. We have effectively taken our replicates into account by collapsing across them within each sample, but this can be dangerous. If there is wide variation across replicates of a particular sample, then taking the mean of three replicates could produce an inaccurate estimate of the ‘true’ sample value. Ideally, we want to find a solution where instead of collapsing this variation, we add it to our statistical model so that we can better understand what amount of variation is randomly coming from within technical replicates, and in turn what amount of variation is actually due to potential differences in our experimental groups.

To achieve this, we need to model both the fixed effect of groups and the random effect of replication across western blot gels. When we use both fixed and random effects, this is referred to as a linear mixed model (LMM). When using this strategy, we find that our effect size remains strong, and our p value is low. But importantly, we now have strong observed power (**Figure 3**). This suggests that we can achieve greater sensitivity in our WB experiment when using this approach. Specifically, if we implement careful counterbalancing while designing our experiments, then we can use the variability between gels to our advantage during analysis using linear mixed effects model(45).

LMM is recommended because it takes into account both the multiple observations within a single subject/animal in a given condition and differences across subjects observed in multiple conditions. This reduces chances of inaccurate p-values and improves reliability(46). Further, treating both subjects and replication as random effects generalizes the results to the population of subjects and also to the population of conditions(47).

### REAL WORLD APPLICATION OF blotRig SOFTWARE FOR WESTERN BLOT EXPERIMENTAL DESIGN, TECHNICAL REPLICATION, and STATISTICAL ANALYSIS

We have designed a user interface that is designed to facilitate appropriate counterbalancing and technical replication for western blot experimental design. Upon starting the blotRig application, the user is prompted to upload a comma separated values (CSV) spreadsheet. This spreadsheet should include separate columns for subject ID and experimental group. The user is then prompted to enter the total number of lanes that are available on their particular western blot gel apparatus. The blotRig software will first run a quality check to confirm that each subject ID (unique sample or subject) is only found in one experimental group. If duplicates are found, a warning will be shown that specifies which subjects are repeated across groups. If no errors are found, a centered gel map will be generated that illustrates the western blot gel lanes into which each subject should be loaded (**Figure 4A**). The decision for each lane loading is based on two main principles outlined above: 1) each western blot gel should hold a representative sample of each experimental group 2) samples from the same experimental group are not loaded in adjacent lanes whenever possible. This ensures that proper counterbalancing is achieved so that we can limit the chances that the inherent variability within and across western blot gels is confounded with the experimental groups that we are interested in experimentally testing.

**Figure 4.**
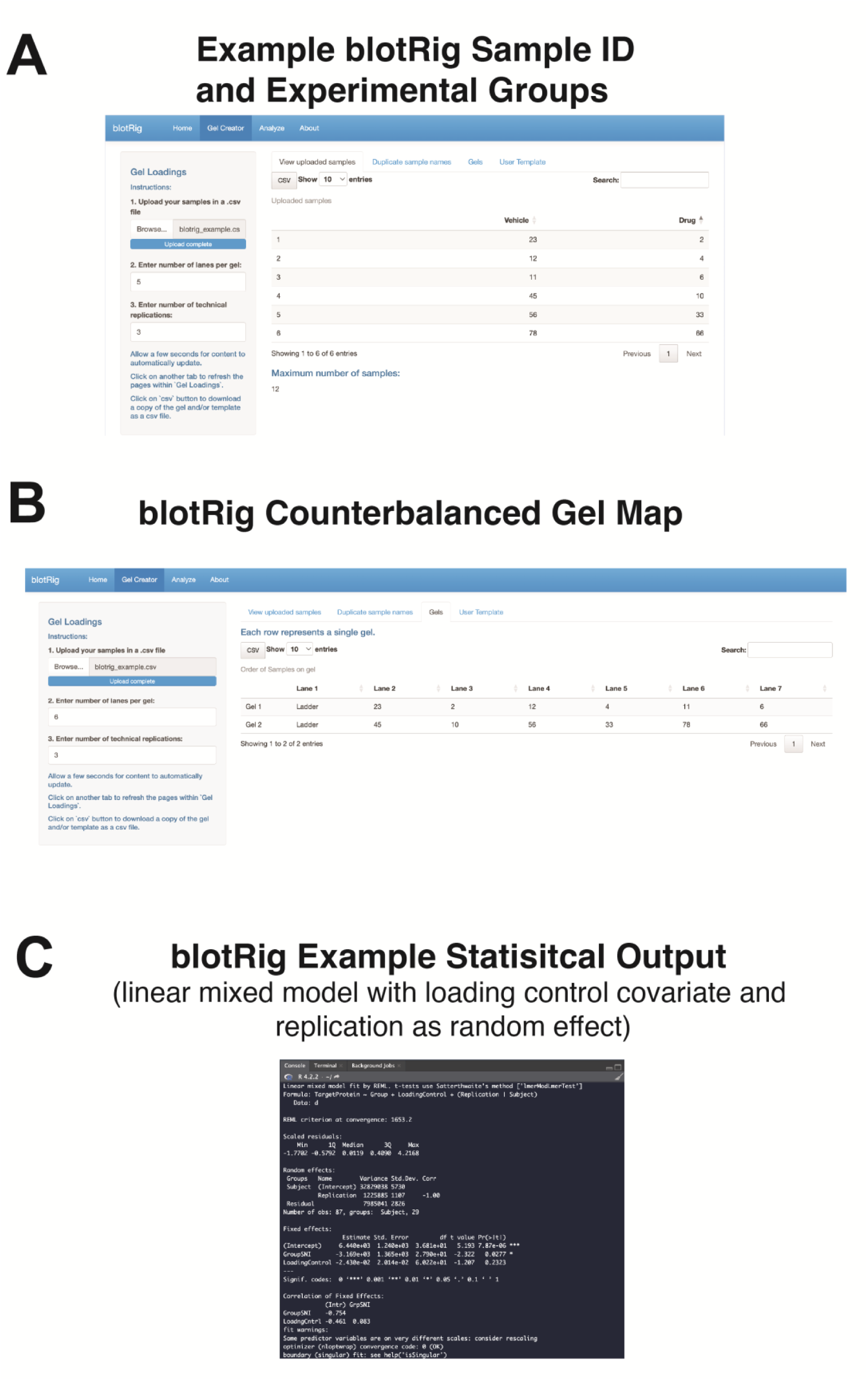
Example of the blotRig interface. **A,** Illustration of the blotRig interface. User has entered their sample IDs, experimental groups, and the number of lanes per western blot gel. **B,** The blotRig system then creates a counterbalanced gel map that ensures each gel contains a representative from each experimental group. This illustration shows the exact lane for each gel in which each sample should be run. **C.** Example output from linear mixed model, indicating random and fixed effects.

Once the gel map has been generated, the user can then select to export this gel map to a CSV spreadsheet. This sheet is designed to clearly show which gel each sample is on, which lane on each gel a sample is found, what experimental group each sample belongs to, and importantly, a repetition of each of these values for three technical replicates (**Figure 4B**). User will also see columns for Target Protein and Loading Control. These are the cells where the user can then input their densitometry values upon completing their western blot runs. Once this spreadsheet is filled out, it is then ready to go for blotRig analysis.

To analyze western blot data, users can upload the completed template that was exported in the blotRig experimental design phase under the ‘Analysis’ tab. The blotRig software will first ask the user to identify which columns from the spreadsheet represent Subject/SampleID, Experimental Group, Protein Target, Loading Control, and Replication. The blotRig software will again run a quality check to confirm that there are no subject/sample IDs that are duplicated across experimental groups. If no errors are found, the data will then be ready to analyze. The blotRig analysis will then be run, using the principles discussed above. Specifically, a linear mixed-model runs using the *lmer* R package, with Experimental Group as a fixed effect, Loading Control as a covariate, and Replication (nested within Subject/Sample ID) as a random factor. Analytical output is then displayed, giving a variety of statistical results from the linear mixed model output table, including fixed and random effects and associated p values (**Figure 4C**). These outputs can be directly reported in the results sections of papers, improve the statistical rigor of published WB reports. In addition, since the entire pipeline is opensource, the blotRig code itself can be reported to support transparency and reproducibility.

## Discussion

Although the western blot technique has proven to be a workhorse for biological research, the need to enhance its reproducibility is critical(11, 17, 26). Current qWB assay methods are still lacking for reproducibly identifying true biological effects(11). We provide a systematic approach to generate quantitative data from western blot experiments that incorporates key technical and statistical recommendations which minimize sources of error and variability throughout the western blot process. First, our study shows that experimenters can improve the reproducibility of western blots by applying the experimental recommendations of determining the linear range for each primary antibody, counterbalancing during experimental design, and running technical triplicates(11, 26). Furthermore, these experimental implementations allow for application of the statistical recommendations of incorporating loading controls as covariates and analyzing gel and subject as random effects(48, 49). Altogether, these enable more rigorous statistical analysis that accounts for more technical variability which can improve the effect size, observed power, and p-value of our experiments and ultimately better identify true biological effects.

Biomedical research has continued to rely on p-values for determining and reporting differences between experimental groups, despite calls to retire the p-value(50). Power (sensitivity) calculations have also become increasingly common. In brief, p-values and the related alpha value are associated with Type I error rate – the probability of rejecting the null hypothesis (i.e., claiming there is an effect) when there is no true effect(51). On the other hand, power effectively controls for the probability of rejecting the null hypothesis (i.e. stating there is not effect) when there is indeed a true underlying effect – a concept that is closely related to reducing the Type II error rate(48, 52). Critically, empirical evidence estimates that the median statistical power of studies in neuroscience is between ∼8% and ∼31%, yet best practices suggest that an experimenter should aim to achieve a power of 80% with an alpha of 0.05(18). By being underpowered, experiments are at higher likelihood of producing a false inference. If an underpowered experiment is seeking to reproduce a previous observation, the resulting false negative may throw into question the original findings and directly exacerbate the reproducibility crisis(48). Even more alarmingly, a low power also increases the likelihood that a statistically significant result is actually a false positive due to small sample size problems (51). In our analyses, we show that our technical and statistical recommendations lowers the p-value (indicates that the observed relationship between variables is less likely to be due to chance) as well as observed power of our experiments. This translates into the ability to better avoid false negatives when there is a true effect as well as reduce the likelihood of false positives when there is not a true experimental effect, both of which will ultimately improve the reproducibility of qWB assay experiments.

Another useful component of statistical analyses that is not as commonly reported but is critically related to p-value and power is effect size. Effect size is a statistical measure that describes the magnitude of the difference between two groups in an experiment(53). It is used to quantify the strength of the relationship between the variables being studied(53). The estimated effect size is important because it answers the most frequent question that researchers ask: how big is the difference between experimental groups, or how strong is the relationship or association?(53). The combination of standardized effect size, p-value and power reflect crucial experimental results that can be broadly understood and compared with findings from other studies(52), thus improving comparability of qWB experiments. In particular, studies with large effect sizes have more power: we are more likely to detect a true positive experimental effect and avoid the false negative if the underlying difference between experimental groups is large(37). In some cases, the calculated effect size is greatly influenced by how sources of variance are handled during analysis(11). Our results demonstrate that by reducing the residual variance (by modeling the random effect of replication) the estimated effect size of our experiment increases. This could mean that the magnitude of the difference between the groups in our experiment is larger than it was originally thought to be. This could be due to a variety of factors such as improving the experimental design, sample size, or the measurement of the variables(11). Likewise, conducting a power analysis is an essential step in experimental design that should be done before collecting data to ensure that the study is adequately powered to detect an effect of a certain size(54).

Increasingly, power analysis is becoming a requirement for publications and grant proposals(55). This is because a study with low statistical power is more likely to produce false negative results, which means that the study may fail to detect a real effect that actually exists. This can lead to the rejection of true hypotheses, wasted resources, and potentially harmful conclusions. In brief, given an experimental effect size and variance, we can calculate the sample size needed to achieve an alpha of 0.05 and power of 0.8; an increased sample size reduces the standard error of mean (SEM), which is the measured spread of sample means and consequently increases the power of the experiment(56). We have demonstrated that our experimental and statistical recommendations lead to a lower p value (Figure 3C) and effect size (Figure 3B) without changing the sample size. This may be of greatest interest to researchers: more rigorous analytics ultimately improves experimental sensitivity without relying solely on increasing the sample size.

Reducing the sample size of an experiment can be beneficial for several reasons, one of which is cost-effectiveness. A smaller sample size can lead to a reduction in the number of animals or other resources that are needed for the study, which can result in lower costs. Additionally, it can also save time and reduce the duration of the experiment, as fewer subjects need to be recruited, and the data collection process can be completed more quickly. However, it is important to note that reducing the sample size can also lead to decreased statistical power. As a result, reducing sample size too much can increase the risk of a type II error, failing to detect significance when there is a true effect(52).Therefore, it is important to consider the trade-off between sample size and power when designing an experiment, and to use statistical techniques like power analysis to ensure that the sample size is sufficient to detect an effect of a certain size.

Moreover, when using animals in research, it’s always important to consider the ethical aspect and the 3Rs principles of reduction, refinement, and replacement(45).

There has been recent recognition that an appropriate study design can be achieved by balancing sample size (n), effect size, and power(29). The experimental and statistical approach presented in this study provide insight into how more rigorous planning for western blot experimental design and corresponding statistical analysis without depending on p-values only can acquire precise data resulting in true biological effects. Using blotRig as a standardized, integrated western blot methodology, quantitative western blot may become highly reproducible, reliable, and a less controversial protein measurement technique(16, 27, 57).

## Abbreviations

qWB: (quantitative western blot)
ELISA: (enzyme linked immunosorbent assay)
SARS-CoV2: (severe acute respiratory syndrome coronavirus 2)
ANCOVA: (analysis of covariance)
ANOVA: (analysis of variance)
SCI: (spinal cord injury)
SNI: (Spared nerve injury)
AMPA: (α-amino-3-hydroxy-5-methyl-4-isoxazoleproprionic acid)
GluA1: (Glutamate receptor 1)
GluA2: (Glutamate receptor 2)
Linear mixed models: (LMM)
(TTBS): Tris-buffered saline containing 0.1% Tween 20
(PAGE): polyacrylamide gel electrophoresis
(SEM): standard error of mean.

## Data availability

The datasets and computer code generated or used in this study are available upon request to Adam Ferguson

## Acknowledgments

The authors are thankful to Dr. Jenny Haefeli and Carlos De Almeida, phD candidate for their helpful contributions.

## Author contributions

A.R.F., J.R.H. and K.M. conceived and designed the experiments; C.O., and J.R.H. performed the experiments; A.R.F., A.C., A.T.E., K.A.F and J.R.H. analyzed the data; C.O., A.C., A.T.E., K.A.F and J.R.H. wrote the paper with input from all other authors.

## Funding and additional information

This work was supported by a National Institutes of Health/National Institute of Neurological Disorders and Stroke grant (R01NS088475) to A. R. F. NIH NINDS: R01NS122888, UH3NS106899, U24NS122732, US Veterans Affairs (VA): I01RX002245, I01RX002787, I50BX005878, Wings for Life Foundation, Craig H. Neilsen Foundation. The content is solely the responsibility of the authors and does not necessarily represent the official views of the National Institutes of Health. Correspondence and requests for materials should be addressed to A.R.F.

## Supporting information

This article contains supporting information. You can access the blotRig analysis software, which includes code for inputting experimental parameters for all Western blot analysis, through the following link: https://github.com/ucsf-ferguson-lab/blotRig/

